# NBI-921352, a First-in-Class, Na_V_1.6 Selective, Sodium Channel Inhibitor That Prevents Seizures in *Scn8a* Gain-of-Function Mice, and Wild-Type Mice and Rats

**DOI:** 10.1101/2021.08.29.458125

**Authors:** JP Johnson, Thilo Focken, Kuldip Khakh, Parisa Karimi Tari, Celine Dube, Aaron Williams, Jean-Christophe Andrez, Girish Bankar, David Bogucki, Kristen Burford, Elaine Chang, Sultan Chowdhury, Richard Dean, Gina de Boer, Shannon Decker, Christoph Dehnhardt, Mandy Feng, Wei Gong, Samuel J Goodchild, Michael Grimwood, Abid Hasan, Angela Hussainkhel, Qi Jia, Stephanie Lee, Jenny Li, Sophia Lin, Andrea Lindgren, Verner Lofstrand, Janette Mezeyova, Rostam Namdari, Karen Nelkenbrecher, Noah Gregory Shuart, Luis Sojo, Shaoyi Sun, Matthew Taron, Matthew Waldbrook, Diana Weeratunge, Steven Wesolowski, Michael Wilson, Zhiwei Xie, Rhena Yoo, Clint Young, Alla Zenova, Wei Zhang, Alison J Cutts, Robin P Sherrington, Simon N Pimstone, Raymond Winquist, Charles J Cohen, James R Empfield

**Affiliations:** Xenon Pharmaceuticals Inc., In Vivo Biology; Xenon Pharmaceuticals Inc., In Vitro Biology; Xenon Pharmaceuticals Inc., Scientific Affairs; Xenon Pharmaceuticals Inc., Chemistry; Xenon Pharmaceuticals Inc., Compound Properties; Xenon Pharmaceuticals Inc., Translational Drug Development; Xenon Pharmaceuticals Inc., Executive Team

## Abstract

NBI-921352 (formerly XEN901) is a novel sodium channel inhibitor designed to specifically target Na_V_1.6 channels. Such a molecule provides a precision-medicine approach to target *SCN8A*-related epilepsy syndromes (*SCN8A*-RES), where gain-of-function (GoF) mutations lead to excess Na_V_1.6 sodium current, or other indications where Na_V_1.6 mediated hyper-excitability contributes to disease (Gardella and Moller, 2019; Johannesen et al., 2019; Veeramah et al., 2012). NBI-921352 is a potent inhibitor of Na_V_1.6 (IC_50_ 0.051 µM), with exquisite selectivity over other sodium channel isoforms (selectivity ratios of 756X for Na_V_1.1, 134X for Na_V_1.2, 276X for Na_V_1.7, and >583X for Na_V_1.3, Na_V_1.4, and Na_V_1.5). NBI-921352 is a state-dependent inhibitor, preferentially inhibiting activated (inactivated or open) channels. The state dependence leads to potent stabilization of inactivation, inhibiting Na_V_1.6 currents, including resurgent and persistent Na_V_1.6 currents, while sparing the closed/rested channels. The isoform-selective profile of NBI-921352 led to a robust inhibition of action-potential firing in glutamatergic excitatory pyramidal neurons, while sparing fast-spiking inhibitory interneurons, where Na_V_1.1 predominates. Oral administration of NBI-921352 prevented electrically induced seizures in a *Scn8a* GoF mouse, as well as in wild-type mouse and rat seizure models. NBI-921352 was effective in preventing seizures at lower brain and plasma concentrations than commonly prescribed sodium channel inhibitor antiseizure medicines (ASMs) carbamazepine, phenytoin, and lacosamide. NBI-921352 was well tolerated at higher multiples of the effective plasma and brain concentrations than those ASMs. NBI-921352 is entering phase II proof-of-concept trials for the treatment of *SCN8A-*developmental epileptic encephalopathy (*SCN8A*-DEE) and adult focal-onset seizures.

## Introduction

Na_V_1.6 voltage-gated sodium channels are widely expressed in the brain and are important contributors to neural excitability (Meisler, 2019; Royeck et al., 2008). Mutations in the *SCN8A* gene result in malfunction of Na_V_1.6 sodium channels and cause a spectrum of *SCN8A*-related syndromes in humans, and disruptions of mouse Na_V_1.6 likewise disrupt normal physiology (Burgess et al., 1995; Gardella and Moller, 2019; Johannesen et al., 2019; Meisler, 2019; Veeramah et al., 2012; Wagnon et al., 2015). Variants of Na_V_1.6 channels can result in either gain or loss of function. Loss-of-function (LoF) variants in humans are generally associated with autism spectrum disorders with cognitive and developmental delay without epilepsy (Inglis et al., 2020; Liu et al., 2019), but, in some cases, can lead to late-onset seizures. In mice, LoF variants of Na_V_1.6 lead to motor impairment but increase seizure resistance (Hawkins et al., 2011; Martin et al., 2007). Gain-of-function (GoF) variants in human *SCN8A* generally result in early-onset *SCN8A*-related epilepsy syndromes (*SCN8A*-RES). The most severe of these epilepsy syndromes is *SCN8A* developmental and epileptic encephalopathy (*SCN8A*-DEE) (Gardella and Moller, 2019; Hammer et al., 2016; Johannesen et al., 2019). Most *SCN8A*-RES patients carry de novo heterozygous missense variants that lead to a gain of function of the Na_V_1.6 channel, though inherited and bi-allelic variants have been reported (Gardella and Moller, 2019; Wengert et al., 2019). *SCN8A*-DEE patients present early in life with seizure onset usually occurring in the first year of life. After seizure onset, patients begin to miss developmental milestones and display additional symptoms, including cognitive and motor delay, hypotonia and cortical blindness. *SCN8A*-DEE individuals are predisposed to early death, including sudden unexplained death in epilepsy (SUDEP). While *SCN8A*-RES patients often have treatment-resistant seizures, many can achieve seizure reduction or seizure freedom upon treatment with anti-seizure medicines (ASMs) that non-selectively inhibit voltage-gated sodium channels, like phenytoin (Boerma et al., 2016; Braakman et al., 2017). *SCN8A*-RES patients may require doses that are higher than those prescribed for most epilepsy patients and, as a result, can be more prone to drug-related adverse events (Boerma et al., 2016; Gardella and Moller, 2019). Even with high doses and multiple ASMs, many patients continue to have uncontrolled seizures as well as extensive comorbidities. The aggressive pharmacotherapy required to protect *SCN8A* patients from life-threatening seizures often comes with attendant side effects that would not be tolerated in less severely impacted populations.

Existing sodium channel inhibitor ASMs are nonselective, blocking all voltage-gated sodium channel isoforms at similar plasma or brain concentrations. This lack of selectivity likely limits the benefits of sodium channel inhibitors since LoF variants of Na_V_1.1 are known to impair inhibitory interneuron function and cause generalized epilepsy with seizures plus (GEFS+) and *SCN1A*-DEE (Dravet Syndrome) (Catterall et al., 2010; Claes et al., 2001; Escayg et al., 2000; Gennaro et al., 2003). Thus, inhibiting Na_V_1.1 may counter the benefit of inhibiting the sodium channels of excitatory neurons. Inhibiting Na_V_1.4 and Na_V_1.5 currents is also undesirable since those channels are critical for facilitating contraction of skeletal and cardiac muscles, respectively (Chen et al., 1998; Ptacek et al., 1991; Rojas et al., 1991).

We hypothesized that a selective inhibitor of Na_V_1.6 could provide a safer and more effective treatment for patients with *SCN8A*-RES and might also be more broadly efficacious in more common forms of epilepsy. An extensive medicinal-chemistry effort produced NBI-921352, the first potent and selective inhibitor of Na_V_1.6 channels (Neurocrine, 2019). We explored the profile of NBI-921352 in vitro, ex vivo and in three preclinical in vivo rodent seizure models, including electrically induced seizure assays in genetically engineered mice bearing heterozygous *Scn8a* GoF Na_V_1.6 channels (N1768D), as well as in wild-type mice and rats.

## Results

### In vitro NaV potency and selectivity

Human Na_V_ channel isoforms hNa_V_1.1, hNa_V_1.2, hNa_V_1.3, hNa_V_1.4, hNa_V_1.5, hNa_V_1.6, and hNa_V_1.7 were heterologously expressed in HEK-293 cells, and the potency and isoform selectivity of NBI-921352 (Figure 1, Table 1) was determined by automated patch-clamp techniques. NBI-921352 potently inhibited hNa_V_1.6 channel currents with an inhibitory concentration 50% (IC_50_) of 0.051 µM (95% CI: 0.030 to 0.073 µM; N=3) calculated from 3 biological replicates. Inhibition of other human Na_V_1.X isoforms required higher concentrations of NBI-921352 with IC_50_’s of 39 µM (95% CI: 31 to 47 µM; N=3) for hNa_V_1.1, 6.9 µM (95% CI: 1.6 to 12 µM; N=3) for hNa_V_1.2, >30 µM for hNa_V_1.3, >30 µM for hNa_V_1.4, >30 µM for hNa_V_1.5, and 14 µM (95% CI: 6.4 to 22 µM; N=3) for hNa_V_1.7. These potencies provide selectivity ratios for hNa_V_1.6 versus the other hNa_V_ isoforms (IC_50_ hNa_V_1.X / IC_50_ hNa_V_1.6) of 756 (Na_V_1.1), 134 (Na_V_1.2), 276 (Na_V_1.7) and >583 (Na_V_1.3, Na_V_1.4, Na_V_1.5).

**Figure 1.**
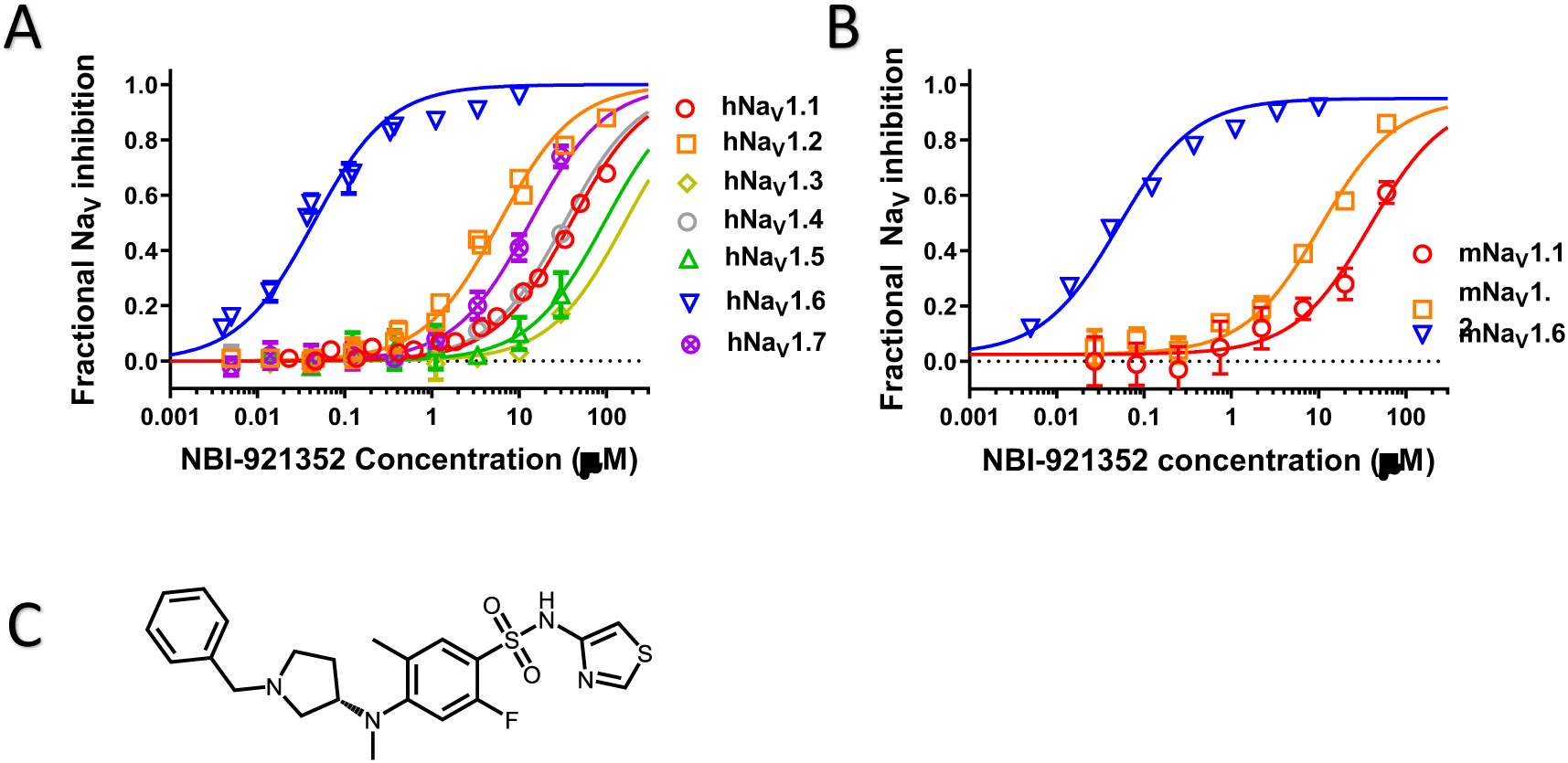
Potency and isoform selectivity of NBI-921352 for human and mouse Na_V_ channels. Concentration-response curves were generated by automated patch-clamp electrophysiology using the Sophion Qube. Concentration-response curves were generated for human (A) or mouse (B) Na_V_ channel isoforms heterologously expressed in HEK293 cells. The analysis included only those cells that met pre-specified acceptance criteria for seal quality, current amplitude, and series resistance. Normalized data from all cell recordings at a concentration were grouped together and plotted with GraphPad Prism 8. Details regarding the number of cells analyzed for each Na_V_ channel and concentration can be found in the source data sheet. Error bars indicating the standard error of the mean fraction were plotted for all points, but, in some cases, they were smaller than the data point symbols and, therefore, not visible. The chemical structure of NBI-921352 is shown (C).

**Table 1.** Potency and isoform selectivity of NBI-921352 for human and mouse Na_V_ channels.

|  | Nav1.6 | Nav1.1 | Nav1.2 | Nav1.3 | Nav1.4 | Nav1.5 | Nav1.7 |
| --- | --- | --- | --- | --- | --- | --- | --- |
| **Human IC<sub>50</sub> (μM)** | 0.051 | 39 | 6.9 | >30 | >30 | >30 | 14 |
| **Human Selectivity**<br>**hNav1.X / hNav1.6** | 1 | 756 | 134 | >583 | >583 | >583 | 276 |
| **Mouse IC<sub>50</sub> (μM)** | 0.058 | 41 | 11 |  |  |  |  |
| **Mouse Selectivity**<br>**mNav1.X / mNav1.6** | 1 | 709 | 191 |  |  |  |  |

Since we intended to evaluate in vivo effects of NBI-921352 in mouse seizure models, we also assessed the potency of NBI-921352 in the mouse Na_V_ isoforms that are most highly expressed in the brain, Na_V_1.6, Na_V_1.1, and Na_V_1.2. The potency and selectivity in mouse Na_V_ channels closely paralleled that seen in the human orthologues with IC_50_’s of 0.058 µM (95% CI: 0.046 to 0.070 µM; N=3) for mNa_V_1.6, 41 µM (95% CI: 30 to 52 µM; N=3) for mNa_V_1.1, and 11 µM (95% CI: 8.2 to 14 µM; N=3) for mNa_V_1.2. Selectivity ratios (IC_50_ mNa_V_1.X / IC_50_ mNa_V_1.6) were 709 (Na_V_1.1), and 191 (Na_V_1.2). These data indicate that NBI-921352 potently inhibits both human and mouse Na_V_1.6 channels, and that it does so at concentrations ≥ 134-fold lower than for any of the other channel isoforms tested.

### NBI-921352 inhibited patient-identified variants of Na_V_1.6 channels

Patients with *SCN8A*-RES carry missense variants in the Na_V_1.6 channel. A great number of variants have been identified, with a range of biophysical defects. Since most variants are de novo, many have been identified in only one or a few patients. For this reason, we determined the effectiveness of NBI-921352 to inhibit 9 patient identified variants spread across the channel (Figure 2, Table 2) (Gardella and Moller, 2019; Wagnon and Meisler, 2015). The 9 variants studied have all been identified in *SCN8A*-RES patients and are in Domains II, III, and IV. Inhibition of the mutant channel constructs was evaluated by automated patch-clamp electrophysiological techniques after transient transfection of the human Na_V_1.6 variant construct of interest into Expi293F™ cells. All the variants were sensitive to inhibition by NBI-921352. Observed IC_50_s for inhibition were 0.051 µM (WT mean from Figure 1), 0.031 µM (95% CI: 0.027 to 0.037 µM) (T767I), 0.021 µM (95% CI: 0.017 to 0.026 µM) (R850Q), 0.032 µM (95% CI: 0.029 to 0.036 µM) (N984K), 0.035 µM (95% CI: 0.029 to 0.043 µM) (I1327V), 0.039 µM (95% CI: 0.031 to 0.050 µM) (N1466K), 0.34 µM (95% CI: 0.26 to 0.44 µM) (R1617Q), 0.055 µM (95% CI: 0.046 to 0.064 µM) (N1768D), 0.068 µM (95% CI: 0.054 to 0.085 µM) (R1872W), and 0.035 µM (95% CI: 0.029 to 0.041 µM) (N1877S). We found that 8 of the 9 variants were inhibited with a potency similar to that of the wild-type channel, with most being slightly more potently inhibited. Only one variant, N1617Q, required markedly higher concentrations of NBI-921352 for inhibition, with an IC_50_ for inhibition 6.6-fold higher than that of the wild-type Na_V_1.6 channel. The reduced potency for N1617Q is consistent with the variant residing in the predicted binding site of NBI-921352 in the domain IV voltage sensor, see discussion.

**Figure 2.**
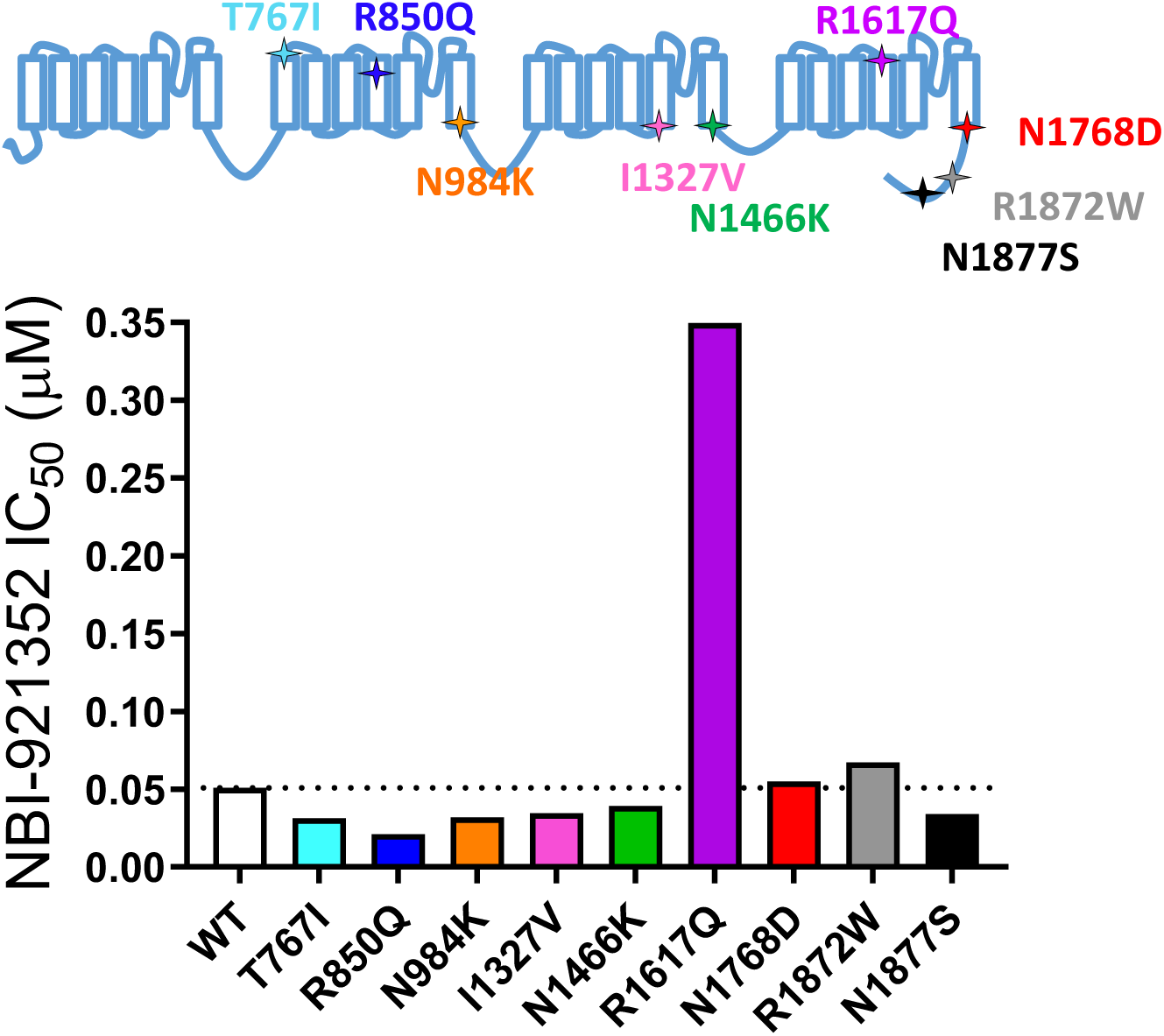
Comparison of NBI-921352 potency on human wild-type Na_V_1.6 and patient-identified variants of Na_V_1.6. IC_50_s were calculated in GraphPad Prism 8 using the Hill equation described in the Materials and Methods. All constructs were transiently transfected into Expi293F™ cells and evaluated by automated patch-clamp electrophysiology using the Sophion Qube. The voltage-clamp methods and data analysis were identical to those used for evaluation of the wild-type channels in Figure 1. Details regarding the number of cells analyzed for each Na_V_ channel and concentration can be found in the source data sheet. The dotted line indicates the IC_50_ for wild-type Na_V_1.6 from Figure 1.

**Table 2.** Comparison of NBI-921352 potency on human wild-type Na_V_1.6 and patient-identified gain-of-function variants of Na_V_1.6.

|  | WT | T767I | R850Q | N984K | I1327V | N1466K | R1617Q | N1768D | R1872W | N1877S |
| --- | --- | --- | --- | --- | --- | --- | --- | --- | --- | --- |
| **hNav1.6 IC<sub>50</sub>**<br>**(μM)** | 0.051 | 0.031 | 0.021 | 0.032 | 0.035 | 0.039 | 0.349 | 0.054 | 0.067 | 0.034 |
| **Fold change**<br>**WT / Variant** | - | 0.6 | 0.4 | 0.6 | 0.7 | 0.8 | 6.8 | 1.1 | 1.3 | 0.7 |

IC<sub>50</sub>s corresponding to Figure 2 are shown in the table and were calculated as indicated for Table 1.

### NBI-921352 is a state-dependent inhibitor

Many small molecule inhibitors of Na_V_ channels bind preferentially to open and or inactivated states (Bean et al., 1983; Courtney et al., 1978; Strichartz, 1976). To query the state dependence of NBI-921352, we measured the apparent potency with two different voltage protocols that favor either the closed (rested) state or activated (open and inactivated) states (Figure 3). Holding the membrane potential at -120 mV induces most channels to reside in the resting state. Brief depolarizations to measure Na_V_1.6 current enabled the determination of an IC_50_ of 36 µM (95% CI: 29 to 47 µM) for rested-state channels. Holding the membrane potential at -45 mV encourages channels to transition into open and inactivated states. Brief hyperpolarizations allow rapid recovery from inactivation for channels that are not bound to drug followed by a short 20 ms test pulse to -20 mV to measure currents from unbound channels (see methods for details). Measuring the ability of NBI-921352 to inhibit reopening of activated channels leads to an apparent IC_50_ of 0.051 µM (Figures 1 and 3). Thus, NBI-921352 strongly prefers activated (open or inactivated) channels, inhibiting them at concentrations more than 750-fold less than those needed to inhibit rested or “peak” sodium currents.

**Figure 3.**
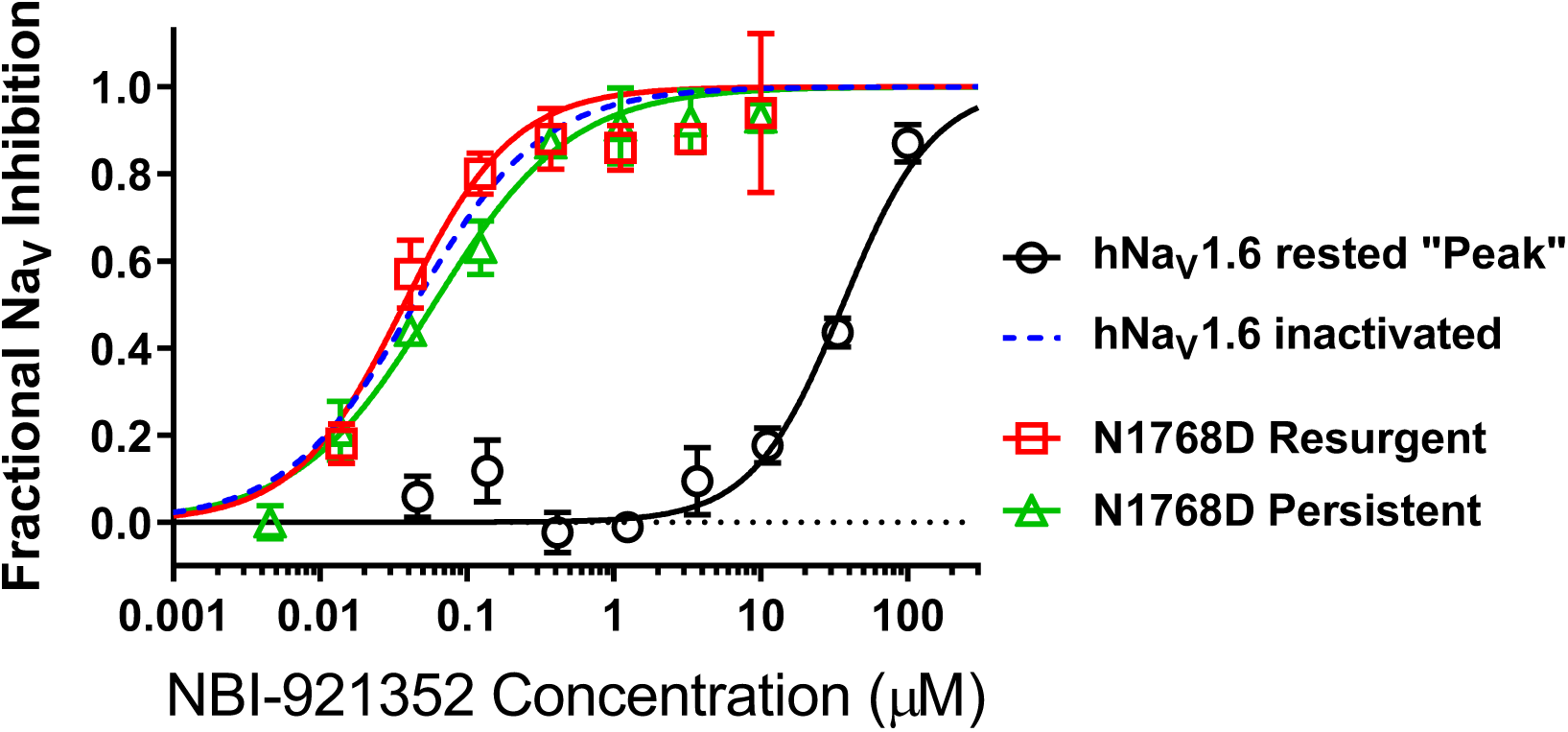
NBI-921352 is a state-dependent inhibitor of Na_V_1.6 and preferentially inhibited activated (open or inactivated) channels. Concentration-response curves were generated for human WT and N1788D channel isoforms heterologously expressed in HEK293 cells. The analysis included only those cells that met pre-specified acceptance criteria for seal quality, current amplitude, and series resistance. Normalized data from all cell recordings at a concentration were grouped together and plotted with GraphPad Prism 8. Details regarding the number of cells analyzed for each Na_V_ channel isoform and concentration can be found in the source data sheet. Error bars indicating the standard error of the mean fraction were plotted for all points. The blue dotted line indicates the concentration-response curve for wild-type Na_V_1.6 from Figure 1. When Na_V_1.6 channels were equilibrated with NBI-921352 at voltages that allow full equilibration with inactivated states (-45 mV), the compound provided potent inhibition, as seen in Figure 1. Measuring persistent or resurgent sodium current after equilibration of cells with NBI-921352 resulted in similar potency (see Materials and Methods and text). Forcing channels to the rested, closed state by hyperpolarizing to -120 mV resulted in very weak inhibition. Current evoked from very negative potentials is sometimes referred to as “peak current”.

### NBI-921352 inhibited persistent and resurgent currents from mutant Na_V_1.6 channels

The state-dependent nature of inhibition is also revealed in other types of voltage-clamp protocols, including those designed to measure persistent or resurgent sodium currents. Some drugs or candidate drugs, like PRAX-330 and Riluzole, have been touted based on their preference for persistent currents, but, in fact, this is a feature of all the compounds in Na_V_ inhibitor class. Apparent differences in persistent current selectivity are driven by differential kinetics and concentration dependences in combination with the electrophysiological protocols chosen for the measurements.

Elevated persistent and or resurgent currents are believed to underlie or contribute to the pathology of many sodium channel related pathologies (Mason et al., 2019; Pan and Cummins, 2020; Potet et al., 2020; Tidball et al., 2020; Zaman et al., 2019). In most conditions, normal Na_V_1.6 channels inactivate rapidly and nearly completely after opening. Persistent currents result from channels that are not stably inactivated – a common phenotype for epilepsy-inducing variants in Na_V_1.6, including N1768D (Tidball et al., 2020; Wagnon et al., 2015). We found that NBI-921352 inhibited N1768D Na_V_1.6 persistent currents (measured as the non-inactivating current 10 ms after initiation of the depolarizing test pulse) with a similar potency as for open and inactivated wild-type Na_V_1.6 channels with an IC_50_ of 0.059 µM (95% CI: 0.044 to 0.082 µM) (Figure 3).

Resurgent currents occur after repolarizing following a strong depolarization as channels redistribute between closed, open, and inactivated states (Raman and Bean, 1997). These resurgent currents are enhanced in many *SCN8A*-RES variants (Pan and Cummins, 2020; Raman et al., 1997). NBI-921352 also effectively inhibited resurgent currents from N1768D channels with apparent IC_50_ of 0.037 µM (95% CI: 0.025 to 0.060 µM).

### NBI-921352 preferentially inhibited excitatory pyramidal neurons and spared inhibitory interneurons

A primary goal of creating Na_V_1.6 selective inhibitors was to spare Na_V_1.1, the voltage-gated sodium channel that is most prevalent in inhibitory interneurons. This should allow the selective targeting of excitatory neurons, where Na_V_1.6 and Na_V_1.2 are believed to be dominant, over inhibitory interneurons. To test this hypothesis, we performed current-clamp experiments in glutamatergic pyramidal neurons from mouse layer 5 neocortex and from fast spiking interneurons in the same region. Application of 0.250 µM NBI-921352 decreased the maximum firing rate in all pyramidal neurons tested (Figure 4A). This reduction was significantly different from control values at current injection intensities between 120 pA and 320 pA (n = 3, p<0.05, paired 2-tailed student’s t-test), except for 310 pA (p = 0.051). In contrast, NBI-921352 subtly increased the number of action potentials (APs) in fast-spiking inhibitory interneurons (Figure 4B) in a statistically significant manner at multiple current injection levels (see Figure 4B). In contrast, carbamazepine inhibited action-potential firing in both pyramidal neurons and in fast-spiking interneurons to a similar degree. The Figure 4 insets show the paired effects of compound on AP number from a stimulus injection of approximately 3X the control Rheobase for that neuron (Pyramidal NBI-921352: 200 pA, Pyramidal carbamazepine: 160 pA, Interneuron NBI-921352: 220 pA, Interneuron carbamazepine: 300 pA ).

**Figure 4.**
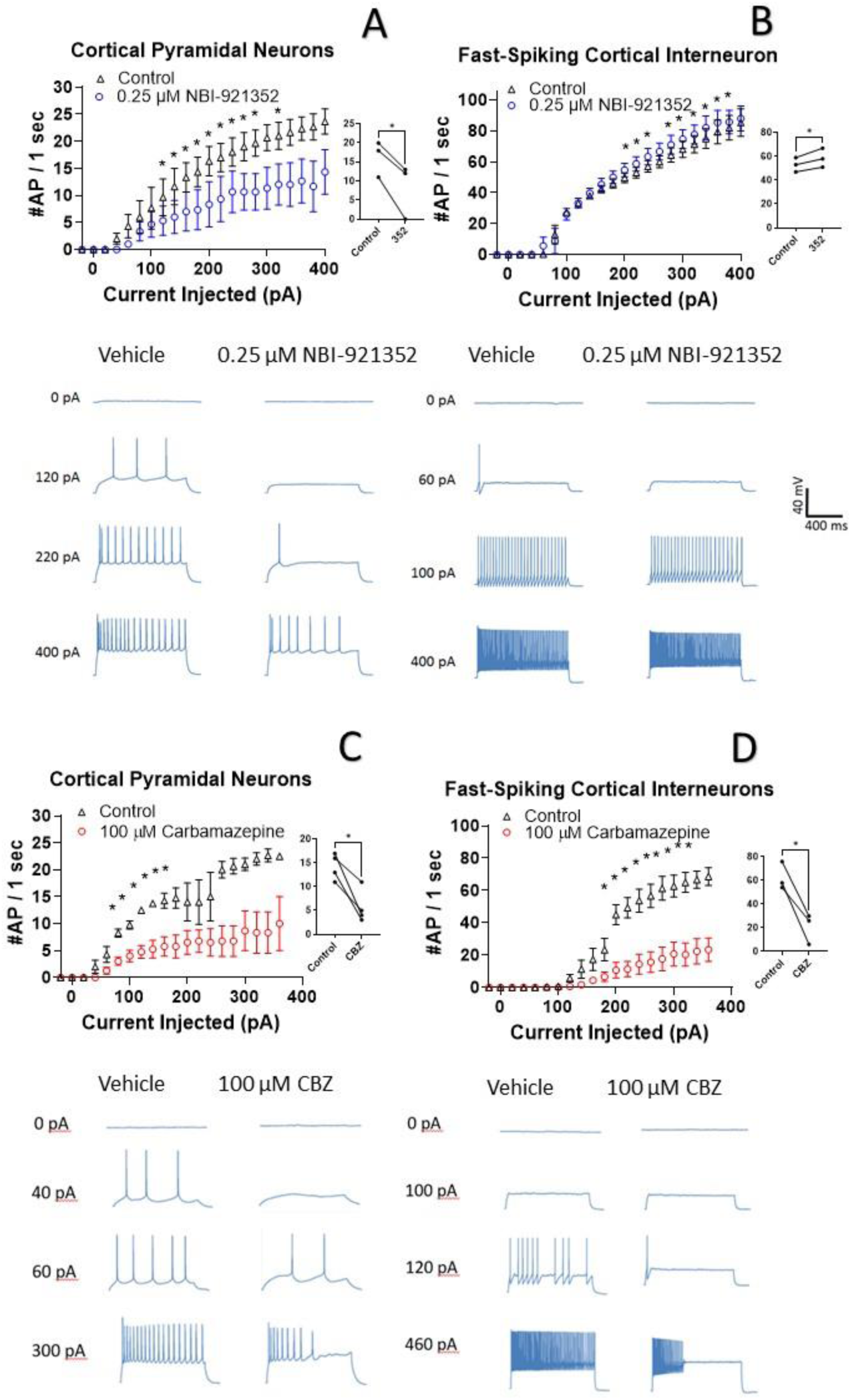
NBI-921352 inhibits firing in pyramidal neurons but spares fast-spiking interneurons. Current input versus action-potential output evaluations in wild-type mouse brain slices treated with vehicle or 0.25 µM NBI-921352 (A & B), or 100 µM carbamazepine (C & D) was plotted. In cortical pyramidal neurons, both NBI-921352 (A) and carbamazepine (C) reduced AP spiking. In fast-spiking cortical interneurons, NBI-921352 increased firing frequency slightly (B), while carbamazepine markedly reduced firing (D). The main upper panels compare average AP count of 3-4 neurons in each condition +/- the standard error of the mean. The inserts in the upper panels show the results for each tested neuron at approximately 3X the cell rheobase. *Indicates a p<0.05 relative to the control condition using a paired, two-way, student’s t-test. The lower panels show recordings for individual representative neurons for each condition. No inhibitors of synaptic inputs were used for these experiments.

### NBI-921352 inhibited electrically induced seizures in *Scn8a^N1768D/+^* mice

A selective inhibitor of Na_V_1.6 should lend itself to the treatment of disease states caused by pathologic gain of function of Na_V_1.6 channels. Hence, we examined the ability of NBI-921352 to inhibit electrically induced seizures in mice with a patient-identified GoF variant in the *Scn8a* gene encoding Na_V_1.6. N1768D is a variant of Na_V_1.6 identified in the first reported *SCN8A*-DEE patient (Veeramah et al., 2012). N1768D Na_V_1.6 channels have impaired voltage-dependent inactivation gating that results in persistent sodium currents and enhanced resurgent currents. Because Na_V_1.6 channels are highly expressed in the neurons of the brain, increased sodium flux in excitatory neurons leads to seizures. Genetically modified mice bearing the same variant (*Scn8a^N1768D/+^*) were created and found to be seizure prone, producing a mouse model with a similar phenotype as that observed in *SCN8A*-DEE patients (Wagnon et al., 2015). Some *Scn8a^N1768D/+^* mice develop spontaneous seizures at age p60 to p100, but seizure onset and frequency is quite variable, making spontaneous seizure studies challenging. In addition, mice rapidly clear NBI-921352, making it extremely difficult to maintain drug plasma and brain levels in an efficacious range for chronic or subchronic dosing experiments. As an alternative means of assessing NBI-921352’s ability to engage Na_V_1.6 channels in vivo, we designed a modified version of the 6Hz psychomotor seizure assay in *Scn8a^N1768D/+^* mice (Barton et al., 2001; Focken et al., 2019). A mild current stimulation (12 mA) evoked robust generalized tonic-clonic seizures (GTC) with hindlimb extension in *Scn8a^N1768D/+^* mice, but not in wild-type littermates.

Oral administration of NBI-921352 two hours prior to electrical stimulation prevented induction of GTC with hindlimb extension in *Scn8a^N1768D/+^* mice in a dose-dependent manner with a 50% effective dose (ED_50_) of 15 mg/kg (95% CI 9.6 to 23 mg/kg, see Figure 5A).

**Figure 5.**
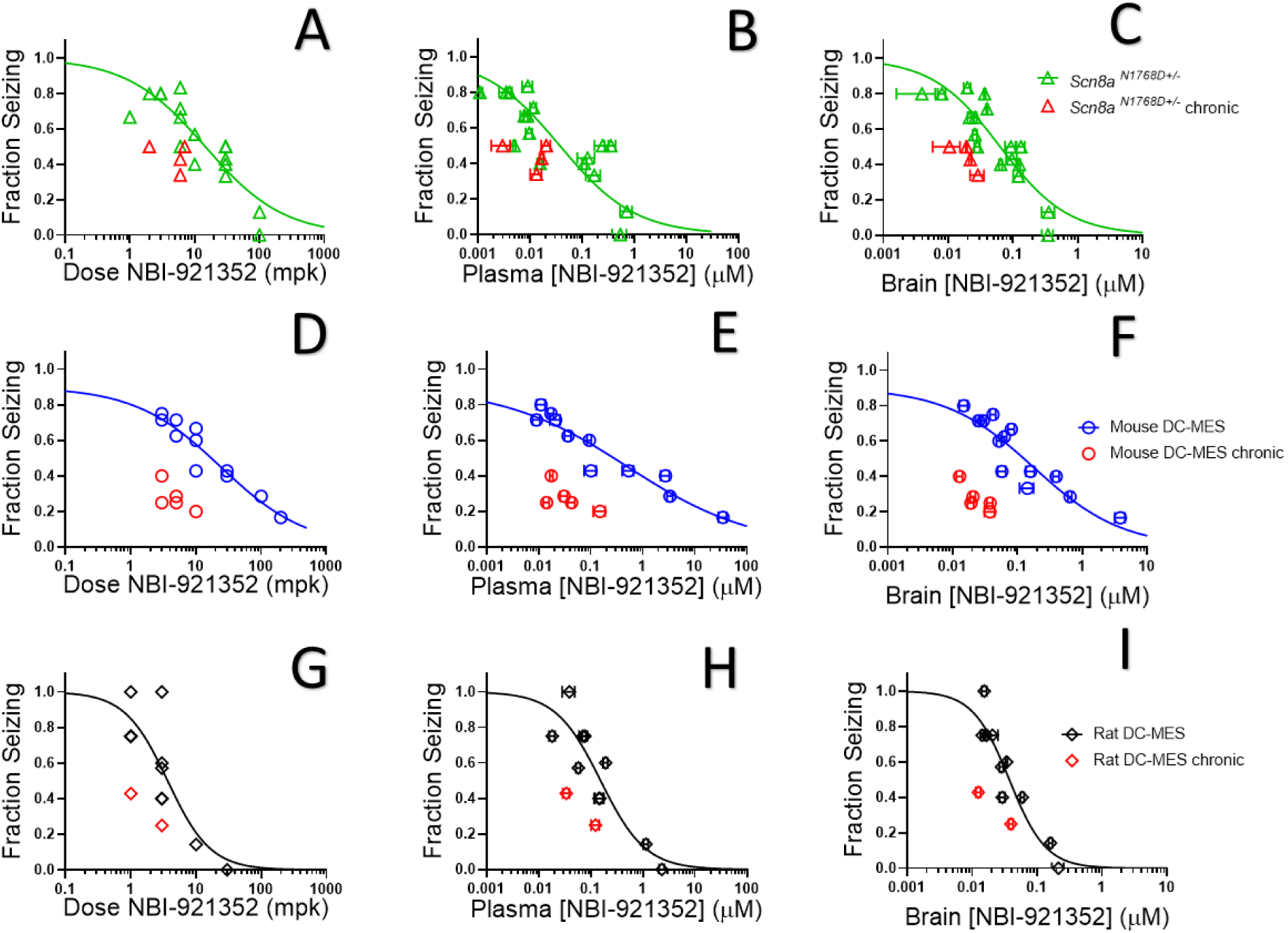
NBI-921352 inhibited electrically induced seizures in rodents. Dose versus efficacy, plasma concentration versus efficacy, and brain concentration versus efficacy relations are shown for *Scn8a*^N1768D+/-^ mice in the modified 6Hz psychomotor seizure assay in A, B, and C, respectively (green open triangles). Dose versus efficacy, plasma concentration versus efficacy, and brain concentration versus efficacy relations are shown for wild-type mice in the DC-MES assay in D, E, and F, respectively (blue open circles). Dose versus efficacy, plasma concentration versus efficacy, and brain concentration versus efficacy relations are shown for wild-type rats in the DC-MES assay in G, H, and I, respectively (black open diamonds). Each point represents the fraction of animals exhibiting a GTC with hindlimb extension after stimulus from a dosing group of 4-8 animals (See source data sheet for all additional information). Horizontal error bars show the standard error of the mean plasma (B, E, H) or brain (C, F, I) concentrations measured from the animals in that dosing group immediately after assay. Where error bars are not visible, they are smaller than the symbols. No error bars are shown for the dose levels (A, D, G), since those were dictated by the experimenter.

After seizure assessment, all animals were euthanized, and the concentration of NBI-921352 was determined in the plasma and brain tissue from each mouse. The average concentrations for each dose group were used to generate plasma concentration and brain concentration versus efficacy relationships (Figures 5B and 5C, respectively). The plasma 50% effective concentration (EC_50_) was 0.037 µM (95% CI 0.018 to 0.090 µM, see Figure 5B). The brain EC_50_ was 0.064 µM (95% CI 0.045 to 0.091 µM, see Figure 5C).

### NBI-921352 inhibited electrically induced seizures in wild-type mice

Na_V_1.6 is an important mediator of neuronal excitability even in animals without GoF mutations. For this reason, we wondered whether NBI-921352 might have broader application in epilepsy beyond *SCN8A*-RES and in other syndromes of neural hyperexcitability. To gain insight into this possibility, we assessed NBI-921352 in a MES assay induced by direct-current electrical stimulus (DC-MES, see methods) in wild-type mice. Figures 5D, E & F show that NBI-921352 prevented GTC with hindlimb extension induction in the DC-MES assay in a dose- and concentration-dependent manner. The ED_50_ for NBI-921352 was 23 mg/kg (95% CI 16 to 34 mg/kg, see Figure 5D). The efficacy of NBI-921352 was also concentration dependent with a plasma EC_50_ of 0.52 µM (95% CI 0.25 to 1.2 µM, see Figure 5E) and a brain EC_50_ of 0.20 µM (95% CI 0.12 to 0.38 µM, see Figure 5F).

### NBI-921352 inhibited electrically induced seizures in wild-type rats

To further explore the preclinical efficacy of NBI-921352, we assessed NBI-921352 in a MES assay induced by direct-current electrical stimulus in wild-type Sprague Dawley rats (see methods). Figures 5G, H & I show that NBI-921352 prevented GTC with hindlimb-extension induction in the rat DC-MES assay in a dose- and concentration-dependent manner. The ED_50_ for NBI-921352 was 3.7 mg/kg (95% CI 2.3 to 7.6 mg/kg, see Figure 5G). The efficacy of NBI-921352 was also concentration dependent with a plasma EC_50_ of 0.15 µM (95% CI 0.09 to 0.31 µM, see Figure 5H) and a brain EC_50_ of 0.037 µM (95% CI 0.028 to 0.054 µM, see Figure 5I).

### Repeated-dosing efficacy in mice and rats

We found that repeated dosing tended to increase efficacy at lower doses and exposures of NBI-921352 than after a single dose (Figure 5, red symbols). Animals were dosed every 12 hours, morning, and evening for 13 doses. Two hours after the 13^th^ dose, on the seventh day, efficacy was tested as in acute-dosing studies. A trend toward improved efficacy was noted in all three assays (see red symbols in Figure 5), but the improvement was not statistically significant when comparing single-dose groups to repeated-dose groups at the same dose level in the same experiment. NBI-921352 did not appreciably accumulate in the plasma or tissue and therefore any trends in improved efficacy were not explained by higher drug concentrations.

### NBI-921352 is effective at lower brain concentrations than three NaV inhibitor ASMs

Both the efficacy and adverse events of Na_V_ inhibitors is driven by the drugs action in the central nervous system (CNS). We found that NBI-921352 was effective in the three preclinical seizure models evaluated at markedly lower brain concentrations than carbamazepine, phenytoin, and lacosamide (Figure 6). The brain EC_50_s for carbamazepine were 9.4 µM, 44 µM, and 36 µM for the *Scn8a^N1768D/+^* 6Hz model, WT mouse DC-MES, and WT rat DC-MES models, respectively. The brain EC_50_s for phenytoin were 18 µM, 13 µM, and 2.6 µM for the *Scn8a^N1768D/+^* 6Hz model, WT mouse DC-MES, and WT rat DC-MES models, respectively. The brain EC_50_s for lacosamide were 3.3 µM, 7.1 µM, and 4.3 µM for the *Scn8a^N1768D/+^* 6Hz model, WT mouse DC-MES, and WT rat DC-MES models, respectively. The lower brain concentrations required for efficacy with NBI-921352 are consistent with the potent inhibition of Na_V_1.6 produced by NBI-921352 (Figure 1).

**Figure 6.**
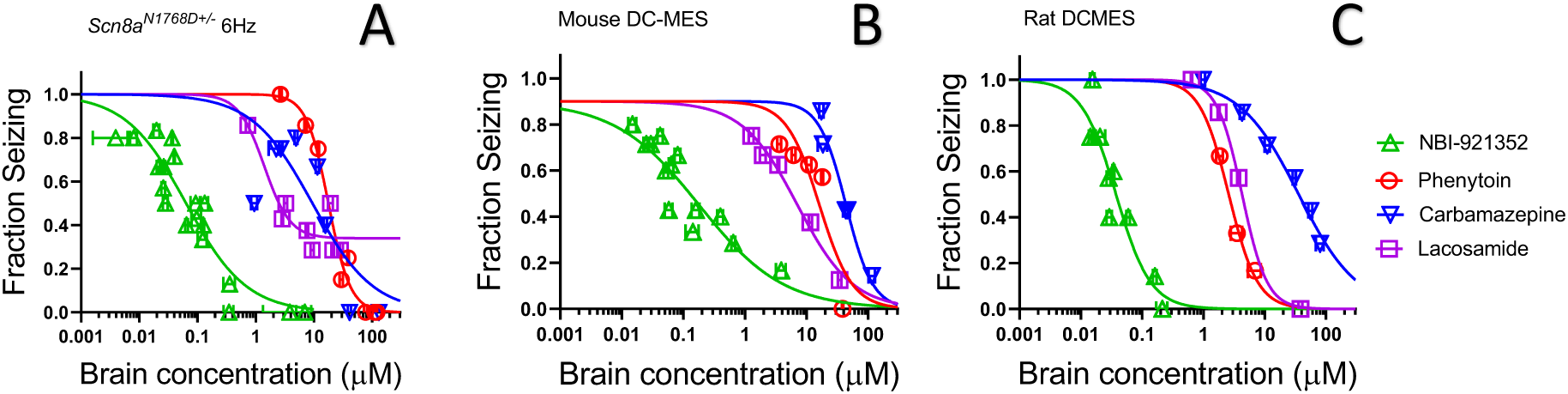
NBI-921352 is more potent than 3 commonly prescribed Na_V_ inhibitor ASMs. Brain concentration versus fraction of animals exhibiting is plotted for NBI-921352 versus that for phenytoin, carbamazepine, and lacosamide in the *Scn8a^N1768D+/-^* modified 6Hz model (A), the wild-type mouse DC-MES model (B), and the wild-type rat DC-MES model (C). Data from animals at a given dose were grouped together and plotted with GraphPad Prism 8. Details regarding the number of animals analyzed per model and per dose group can be found in the source data sheet. Error bars indicating the standard error of the mean concentration were plotted for all points on the concentration-response curves.

### NBI-921352 provided improved separation between efficacy in rats and behavioral signs

The intent of creating a highly selective Na_V_1.6 antagonist was to reproduce or improve on the efficacy of classic, nonselective, sodium channel inhibitor drugs while reducing or preventing the adverse events caused by polypharmacy with other sodium channel and non-sodium channel targets. If sparing Na_V_1.1 and other off-target interactions does, in fact, reduce adverse events, then even higher receptor occupancy of Na_V_1.6 might be achievable, thereby further improving efficacy.

To evaluate our hypothesis, we compared the window between the plasma concentrations required for efficacy (plasma EC_50_) relative to the minimal plasma concentration at which treated rats showed behavioral signs of adverse effects as reported by the blinded experimenter (Figure 7). We made this comparison both for NBI-921352 and for several widely used Na_V_ inhibitor ASMs: carbamazepine, phenytoin, and lacosamide.

**Figure 7.**
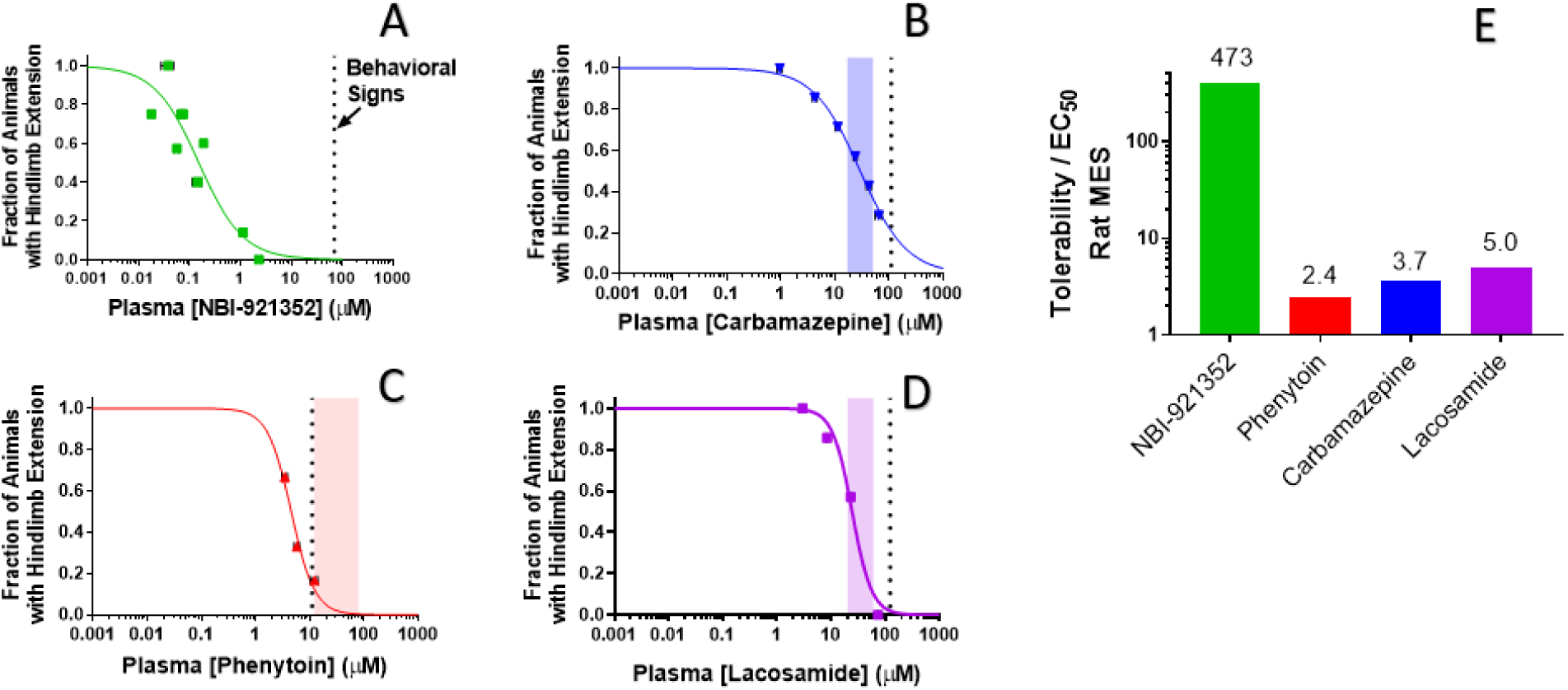
Rat efficacy compared to acute tolerability for NBI-921352 relative to Na_V_ inhibitor ASMs. Plasma concentration versus efficacy data is shown for the rat DC-MES assay for NBI-921352 (A), carbamazepine (B), phenytoin (C), and lacosamide (D). The vertical dotted lines indicate the lowest plasma concentration at which a rat was observed to exhibit atypical behavioral signs indicative of an adverse reaction to drug in the assay format. Animals exhibiting such signs were excluded from efficacy evaluation. The shaded bars in B, C, and D indicate the approximate human plasma concentrations observed in clinical practice. Data from animals at a given dose were grouped together and plotted with GraphPad Prism 8. Details regarding the number of animals analyzed per model and per dose group can be found in the source data sheet. Error bars indicating the standard error of the mean concentration were plotted for all points on the concentration-response curves. Panel E shows the ratio of the (rat plasma EC_50_ / the plasma concentration where behavioral signs were noted for each compound).

NBI-921352 was well tolerated in these studies up to a plasma concentration of 71 µM. Dividing this concentration by the plasma EC_50_ of 0.15 µM in the rat DC-MES study results in a behavioral signs concentration (BSC) / plasma EC_50_ ratio of 473-fold. The same calculation was repeated for the established ASMs. The minimal plasma concentrations provoking behavioral signs for carbamazepine, phenytoin, and lacosamide were 110 µM, 11 µM, and 123 µM, respectively. Their plasma EC_50_s were 30 µM, 4.5 µM, and 24.4 µM, respectively. Figure 7E shows BSC / Plasma EC_50_ ratios for carbamazepine (3.7-fold), phenytoin (2.4-fold), and lacosamide (5.0-fold). This data indicates that increasing Na_V_1.6 selectivity can improve the tolerability of Na_V_ inhibitors in rodent-seizure models.

## Discussion

Sodium channel inhibitors have long been, and remain, a mainstay of pharmacotherapy for epilepsy, as well as for pain and other neurologic, cardiac, and skeletal muscle disorders. The diverse range of indications and systems affected by these drugs is a testament to their critical biological role in cellular excitability. A fundamental challenge for these currently marketed sodium channel drugs is that none of them are selective amongst the nine sodium channel isoforms. As a result, drugs targeting the sodium channels of the brain for epilepsy can inhibit both excitatory and inhibitory neurons, limiting their ability to restore balanced neuronal firing.

Reducing Na_V_1.1 current is proconvulsant due to the predominance of Na_V_1.1 in inhibitory interneurons (Catterall et al., 2010; Claes et al., 2001; Escayg et al., 2000; Mistry et al., 2014; Yu et al., 2006), while the opposite is true for Na_V_1.2 and Na_V_1.6 currents (Ben-Shalom et al., 2017; Hawkins et al., 2011; Martin et al., 2007). Na_V_1.2 and Na_V_1.6 are more highly expressed in excitatory neurons (Catterall et al., 2010; Du et al., 2020; Encinas et al., 2020).

Additionally, these nonselective agents can block the channels associated with skeletal muscle (Na_V_1.4), cardiac tissue (Na_V_1.5), and peripheral neurons (Na_V_1.7, Na_V_1.8, Na_V_1.9). Inhibiting these off-target channels can compromise muscular, cardiovascular, and sensory function. These risks are highlighted by the FDA’s recent drug-safety communication for the nonselective Na_V_ inhibitor ASM lamotrigine. Lamotrigine has been linked to cardiac liabilities as a consequence of Na_V_1.5 inhibition (FDA, 2021). Likewise, Na_V_ inhibitors intended as local anesthetics for trigeminal neuralgia or other pain syndromes and class I cardiac antiarrhythmic drugs are often dose limited by CNS adverse events like dizziness, sedation, and cognitive or motor impairment caused by inhibition of central nervous system Na_V_ channels (Caron & Libersa, 1997).

An obvious solution to this isoform selectivity challenge is to pursue a precision-medicine approach and create selective pharmacologic agents that preferentially target the sodium channels specific to the desired target tissue or cell type. This selective approach has been pursued for multiple channel isoforms - particularly for the peripheral Na_V_s associated with pain (Na_V_1.3, Na_V_1.7, and Na_V_1.8). This approach has proven challenging because the nine isoforms of sodium channels, Na_V_1.1-Na_V_1.9, share a high degree of primary and tertiary structural homology. Achieving selectivity with compounds that have tractable pharmaceutical properties has been particularly difficult.

Previous attempts to optimize Na_V_ inhibitors for epilepsy have focused on either drug properties or channel-state dependence. To our knowledge, this is the first description of a centrally penetrant, isoform-selective Na_V_ inhibitor for use in CNS indications, including epilepsy. NBI-921352 represents the first selective inhibitor of Na_V_1.6 that is suitable for systemic oral administration.

The Na_V_1.6 selective profile of NBI-921352 was designed to inhibit activity in excitatory neurons while sparing firing in the inhibitory interneurons where Na_V_1.1 is preferentially expressed. We found that NBI-921352 did, in fact, reduce firing in cortical excitatory pyramidal cells. In contrast, inhibitory interneuron firing was not impaired and was seen to increase slightly. The reason for an increase in interneuron action-potential firing is unclear. We propose that this may be a consequence of network effects that arise from interactions with other neurons synapsed onto the target neurons in the experiment. These data confirm that selective Na_V_1.6 inhibitors can distinguish neuronal subtypes in a way that nonselective inhibitors, like carbamazepine, cannot.

*SCN8A*-RES patients most often carry de novo genetic variants. While some variants are known to be recurrent, many variants are represented by a single patient (Meisler, 2019). We therefore wanted to assure that NBI-921352 inhibition was not limited to wild-type Na_V_1.6 channels. We tested 9 distinct, patient-identified variants and found that 8 of them were inhibited by NBI-921352 at very similar concentrations as wild-type channels (Figure 2). One variant, R1617Q, was found to be 6.8-fold less sensitive to inhibition than the wild-type channel. R1617Q has been identified in multiple *SCN8A*-DEE patients and is in the domain IV voltage-sensor domain (VSD4). NBI-921352 is an aryl sulfonamide with structural similarity to the Na_V_1.7 targeted aryl sulfonamides where the binding site has been identified as the Na_V_1.7 VSD4 (Ahuja et al., 2015; McCormack et al., 2013). It is likely that the R1617Q directly or allosterically impairs the tight association of NBI-921252 with Na_V_1.6 due to its proximity to the binding site. Despite this, NBI-921352 remains markedly more potent than existing Na_V_ inhibitor drugs on the N1617Q variant Na_V_1.6 channel. This would suggest that while NBI-921352 may be an effective treatment for *SCN8A*-DEE patients carrying R1617Q variants, higher plasma levels of the compound could be required for efficacy in those patients.

Many *SCN8A*-RES associated variants produce their GoF effects by disrupting or destabilizing the inactivation-gating machinery of Na_V_1.6 channels. This can lead to pathological persistent or resurgent currents that contribute to neuronal hyperexcitability (Pan and Cummins, 2020).

Most known small molecule inhibitors of Na_V_ channels, except some marine toxins like tetrodotoxin, bind preferentially to inactivated or open gating states of the channels and stabilize the channels in inactivated, non-conductive conformations. This state dependence is manifested as a protocol dependence of the apparent drug potency. State dependence inhibition has been described in many ways. Use-dependent, frequency-dependent, resurgent current -selective, and persistent current-selective inhibition are all consequences of a preference for binding to inactivated and/or open channels. Stabilizing inactivated states of the channel reduces persistent and resurgent currents, and this feature has been suggested to contribute to the efficacy of many Na_V_ targeted ASMs including phenytoin, carbamazepine, oxcarbazepine, lacosamide, cannabidiol, and lamotrigine (Wengert and Patel, 2021). NBI-921352 is also highly state dependent, with a >750-fold preference for open and inactivated channels vs. rested, closed-state channels (sometimes referred to as *peak current*). Forcing all Na_V_1.6 channels into the closed state by applying voltages more hyperpolarized than physiological (-140 mV) results in very weak inhibition of the channels (Figure 3). Biasing the channels toward activated states (open or inactivated states) by holding the membrane potential more positive in a protocol designed to monitor inactivated state, resurgent current, or persistent current protocol yields potent inhibition. In physiologic conditions, channels are distributed among closed, open, and inactivated states, thus allowing equilibration of potent inhibition of the channel by NBI-921352.

Increasing the selectivity of a Na_V_ inhibitor provides the expectation of an improved safety profile by reducing adverse events caused by off-target activity. An inherent risk of this approach is the potential loss of efficacy that could come from reduced polypharmacy. We have developed a potent, highly selective Na_V_1.6 inhibitor in NBI-921352. Our studies with NBI-921352 indicate that a Na_V_1.6 specific compound can retain a robust ability to prevent seizures in rodent models at modest plasma and brain concentrations, consistent with the important role of Na_V_1.6 in seizure pathways. Our data also suggests that this selectivity profile does improve the tolerability of NBI-921352 relative to commonly employed nonselective sodium channel ASMs in rodents. Whether these results will translate to humans is not yet established, but Phase I clinical trials have shown that NBI-921352 was well tolerated at plasma concentrations higher than were required for efficacy in the preclinical rodent studies described here. NBI-921352 is currently being developed for both *SCN8A*-DEE epilepsy and adult focal-onset seizures by Neurocrine Biosciences (Neurocrine, 2019). Phase II clinical trials will soon evaluate the efficacy of NBI-921352 in patients (Neurocrine, 2021). These clinical trials will provide the first evidence for whether the robust efficacy and tolerability demonstrated in rodents translates to human epilepsy patients.

## Materials and Methods

### Electrophysiological determination of potency and selectivity

#### Cell lines

Electrophysiology experiments were performed with HEK293 cells either stably transfected or transiently transfected. The stable cell lines were transfected with an expression vector containing the full-length cDNA coding for specific human and mouse sodium channel α-subunit, grown in culture media containing 10% fetal bovine serum, and 0.5 mg/mL Geneticin (G418) at 37°C with 5% CO_2_. The Na_V_1.x stable cell lines and accessory constructs used correspond to the following GenBank accession numbers: Human Na_V_1.1 (NM_006920); mouse Na_V_1.1 (NM_018733.2); human Na_V_1.2 (NM_021007); mouse Na_V_1.2 (NP_001092768.1); human Na_V_1.5 (NM_198056); human Na_V_1.6 (NM_014191); mouse Na_V_1.6 (NM_001077499); human Na_V_1.7 (NM_002977); human Na_V_1.4 (NM_000334); human Na_V_1.3 (NM_0069220). The human Na_V_ β1 subunit (NM_199037) was co-expressed in all cell lines. Human and mouse Na_V_1.6 channels were also coexpressed with human FHF2B (NM_033642) to increase functional expression. Human Na_V_1.2 channels were also coexpressed with Contactin 1 (NM_001843) to increase functional expression.

For studies of mutant channels, cDNA plasmids in pcDNA™4/TO Mammalian Expression Vector were transiently transfected into Expi293F™ cells stably expressing human FHF2b and human SCN1B subunit (polyclonal) background using ExpiFectamine™ 293 Transfection Kits (Gibco,Thermo Fisher Scientific CAT #: A14524). Induction was achieved using Tetracycline (Sigma Aldrich). Transfected cells were used in automated patch-clamp experiments 24 hours postinduction.

#### Na_V_ channel automated planar patch-clamp assay

NBI-921352 requires several seconds to equilibrate with activated channels, and this property of the compound must be taken into consideration in the design of state-dependent assay voltage-clamp protocols.

Data was collected using the Qube 384 (Sophion) automated voltage-clamp platform using single hole plates. To measure inactivated state inhibition, the membrane potential was maintained at a voltage where inactivation is complete. For each Na_V_ channel subtype, the V_h_ used to quantify compound inhibition were as follows: Na_V_1.6 (-45 mV), Na_V_1.1 (-45 mV), Na_V_1.2 (-45 mV), Na_V_1.3 (-45 mV), Na_V_1.5 (- 60 mV), Na_V_1.7 (-60 mV), Na_V_1.4 (-45 mV). The mutant channel hNa_V_1.6^N1768D^ was found to have accelerated run-down compared with wild-type hNa_V_1.6, so the holding potential was adjusted to -60 mV to provide sufficient signal window. The voltage was briefly repolarized to a negative voltage (-150 mV) for 20 milliseconds for (Na_V_1.5, Na_V_1.7, Na_V_1.3, Na_V_1.4) or for 60 milliseconds (for Na_V_1.1, Na_V_1.2, and Na_V_1.6) to allow recovery from fast inactivation, followed by a test pulse to -20 or 0 mV for 10 milliseconds to quantify the compound inhibition. The repolarization step allows compound-free channels to recover from fast inactivation, but compound-bound channels remain inhibited during the subsequent test step. For rested state “Peak” current V_h_ was set to -120 mV. Appropriate filters for minimum seal resistance were applied (typically >500 MΩ membrane resistance), and series resistance was compensated at 100%. The pulse protocols were run at 1 Hz for hNa_V_1.7, hNa_V_1.5, hNa_V_1.3, and hNa_V_1.4 or 0.04 Hz for Na_V_1.6, Na_V_1.1 and Na_V_1.2.

To construct concentration response curves, baseline currents were established after 20 minutes in vehicle (0.5% DMSO). Full inhibition response amplitudes were determined by adding tetrodotoxin (TTX, 300 nM) or tetracaine for Na_V_1.5 (10 µM) to each well at the end of the experiment. Compounds were then exposed at a single concentration for 20 minutes. One-sixth of every experimental plate was dedicated to vehicle-only wells that enabled correction for nonspecific drift (i.e., rundown) of the signal in each experiment. For all channel subtypes, inhibition by the compound reached steady state within 20 minutes of incubation. The current inhibition values (I_(CPD)_) were normalized to both the vehicle (I_control_) and the full response defined by supramaximal TTX (I_TTX_) or tetracaine (for Na_V_1.5) addition responses according to Equation 1:

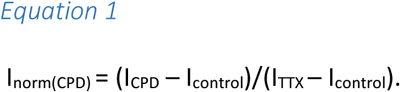

This normalized inhibition was then further normalized to the span of the assay to account for the run-down seen in cells exposed to vehicle alone for 20 minutes as follows:

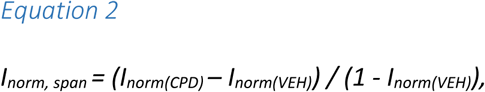

where:

*I_norm, span_* = the current response normalized to within the span of the assay.

*I_norm(CPD)_* = the normalized response in the presence of compound.

*I_norm(VEH)_)* = the normalized response in the absence of compound.

This normalization ensures that the data ranges were between 0 and 1, and there is no rundown in the plots. The normalized data from all cell recordings at a concentration were grouped together and plotted with GraphPad Prism 8, and IC_50_ values were calculated for grouped data using the following version of the Hill equation:

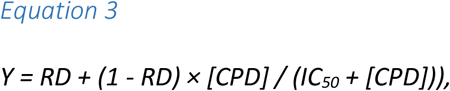

where:

*Y* = the fraction of sodium current blocked in the presence of the compound.

*[CPD]* = the concentration of compound.

*IC_50_* = the IC_50_ concentration.

*RD* = the “rundown” of sodium current in vehicle alone, which is equal to 0 in this case, as the inhibition has already been normalized to the span.

The Hill slope was fixed to 1.

The 95% CI for the IC_50_ from the fitted curve to the mean data were reported unless otherwise noted.

To evaluate inhibition of hNa_V_1.6(N1768D) resurgent currents, synthetic Na_V_β4 peptide (KKLITFILKKTREKKKECLV) was added to the intracellular recording solution at 200 μM and a dedicated protocol to elicit resurgent currents was employed (Barbosa et al., 2015). Cells were voltage clamped at (V_h_ = -80 mV) and subjected to a strong depolarization (+60 mV) for 20 milliseconds. Following the strong depolarization, cells were partially repolarized to the voltage where resurgent current was maximal (-20 mV) for 50 milliseconds, and resurgent current amplitude was measured. This resurgent current-specific waveform was repeated at 5 Hz for 100 s in vehicle, followed by 100 s in test compound, then 300 nM TTX. Fractional inhibition was calculated using the same normalization procedure as above.

Experiments were all performed at 27°C ± 2°C.

#### Automated patch-clamp recording solutions

The recording solutions for Na_V_1.1, Na_V_1.2, Na_V_1.3, Na_V_1.4 and Na_V_1.6 cell line studies contained: Intracellular solution (ICS): 5 mM NaCl, 10 mM CsCl, 120 mM CsF, 0.1 mM CaCl_2_, 2 mM MgCl_2_, 10 mM HEPES (4-(2-hydroxyethyl)-1-piperazineethanesulfonic acid buffer), 10 mM EGTA (ethylene glycol tetraacetic acid); adjusted to pH 7.2 with CsOH. Extracellular solution (ECS): 140 mM NaCl, 5 mM KCl, 2 mM CaCl_2_, 1 mM MgCl_2_, 10 mM HEPES; adjusted to pH 7.4 with NaOH. Solutions with a reversed Na^+^ gradient were used for Na_V_1.5 and Na_V_1.7 studies since they improved technical success. ICS: 120 mM NaF, 10 mM CsCl, 0.1 mM CaCl_2_, 2 mM MgCl_2_, 10 mM HEPES, 10 mM EGTA; adjusted to pH 7.2 with CsOH. ECS: 1 mM NaCl, 139 mM CholineCl, 5 mM KCl, 2 mM CaCl_2_, 1 mM MgCl_2_, 10 mM HEPES; adjusted to pH 7.4 with NaOH. Osmolarity in all ICS and ECS solutions was adjusted with glucose to 300 mOsm/kg and 310 mOsm/kg, respectively.

### Current-clamp recording of cortical pyramidal neurons and inhibitory interneurons

#### Slice preparation

Parasagittal cortical brain slices were prepared from >P21 mice using standard procedures (adapted from Tai et al., PNAS 2014). Briefly, the mouse was deeply anaesthetized with isoflurane and decapitated. The brain was removed and placed into chilled artificial cerebrospinal fluid (aCSF) solution containing (in mM): 125 NaCl, 25 NaHCO3, 2.5 KCl, 1.25 NaH_2_PO_4_, 2 CaCl_2_, 2 MgCl_2_, 10 d-glucose, pH 7.3, osmolarity adjusted to ∼306 mOsm using sucrose. All solutions were saturated with 95% O_2_ and 5% CO_2_ constantly perfused with 95% O_2_/5% CO_2_. Slices with a thickness of 400 µm were prepared using a vibratome (Ted Pella, Inc.). Following sectioning, the slices were placed in a holding chamber and incubated in a water bath at 34°C for 15 minutes. The brain slices were removed from the water bath and held at room temperature for 60 minutes prior to recording.

#### Brain slice electrophysiology assay

All experiments involving rodent subjects were performed in accordance with the guidelines of the Canadian Council on Animal Care (CCAC). Following a 60-minute incubation at room temperature, a brain slice was selected and placed on the stage of an upright microscope (SliceScope Pro 2000, Scientifica). The slice was constantly perfused with room temperature aCSF, containing 0.1% DMSO as a vehicle control, and oxygenated with 95% O_2_/5% CO_2_.The slice was visualized using brightfield microscopy, and a healthy neuron was selected from neocortical layer 5. Whole-cell configuration was achieved with a pipette (bath resistance 4 – 6 MΩ) containing internal solution. Stimulation was applied in current-clamp mode, and consisted of a series of 1000 ms square pulses, beginning at -20 pA and increasing by +20 pA increments (3000 ms between pulses).

Once the recordings in vehicle were completed, and while still holding the patch on the same neuron, the bath solution was changed from 0.1% DMSO in aCSF to 0.25 µM NBI-921352 or 100 µM Carbamazepine in aCSF. The slice was incubated in circulating compound for 10 minutes before repeating the series of square pulse stimulations. Working stock solutions were prepared in DMSO at a concentration of 20 mM.

All data analysis was done offline using ClampFit 10.7 (Molecular Devices). Data are presented as a mean ± SEM. For each sweep, the number of evoked APs was counted, and plotted as a function of current injection (beginning with -20 pA). These generated “input/output” (or “F/I”) curves demonstrating the relationship between stimulus and average AP frequency. Statistical significance was assessed using paired, two-way, student’s t-test applied at each current injection level with significance considered P<0.05.

### Formulation and oral dosing of NBI-921352

#### Vehicle preparation

The vehicle for oral dosing solutions was 0.5% methyl cellulose and 0.2% Tween-80 in deionized (DI) water. DI water (0.8 L) was heated up to 70°C to 80°C. Five grams of methyl cellulose was slowly added to heated DI water. The mixture was stirred until it formed a homogeneous milky suspension. The suspension was moved to a cold room and stirred overnight to get a clear solution. Two milliliters of Tween-80 was added to the clear solution and diluted up to 1 L with DI water. The vehicle solution was stored at 2°C to 8°C.

#### Drug formulation

NBI-921352 was weighed into vials. An appropriate amount of vehicle was added to the NBI-921352 powder then mixed on a T18 ULTRA TURRAX homogenizer (IKA, Wilmington, NC) to create a uniform suspension at the desired concentration. The vials were then wrapped in aluminum foil to protect them from light and placed on a stir plate until the time of dosing. Carbamazepine and lacosamide were formulated in the same manner. Phenytoin was formulated in 0.9% physiological saline.

#### Dosing

NBI-921352, carbamazepine, and lacosamide were administered orally using a stainless-steel gavage needle at a dose volume of 10 ml/kg. Phenytoin was formulated in physiologic saline and was administered intraperitoneally (i.p.) using a 25-gauge needle at a dose volume of 10 mL/kg. All compounds were administered 2 hours prior to electrical seizure induction for all seizure models employed in this study.

### Bioanalytical assessment of plasma and brain concentrations

#### Sample collection

Approximately 0.5 mL of blood was collected from each mouse at the end of the assay via cardiac puncture under deep anesthesia. The blood samples were collected in a syringe and transferred to tubes containing EDTA. Blood was stored at 4°C until centrifuged within 30 minutes of collection. Plasma was harvested and placed on dry ice and stored in a freezer set to maintain a temperature of -70°C to -80°C until analysis. Brains were harvested immediately after blood collection and placed on dry ice prior to storage in a freezer set to maintain a temperature of -70°C to -80°C until analysis.

#### Plasma samples

Extraction of plasma samples was carried out by protein precipitation using acetonitrile. Plasma samples (50 µL) were mixed with 50 µL of internal standard (IS) solution in water followed by addition of 10 µL of concentrated ortho-phosphoric acid and 200 µL of acetonitrile. Samples were vortexed for 30 seconds, centrifuged at 13,000 rpm for 20 minutes, decanted in to a 96-well plate, and further centrifuged at 4,000 rpm for 20 minutes. The samples were analyzed by UHPLC-ESI-MS/MS as described below.

#### Brain samples

Prior to extraction, pre-weighed whole brains were homogenized in 1:1 acetonitrile/water (v/v) (4 mL per mouse brain) using an IKA T18 ULTRA-TURRAX Homogenizer at the setting of 4 for approximately 2 min. The homogenate was centrifuged at 13,000 rpm for 20 min and 50 µL of the supernatant were treated exactly as described above for plasma samples. 50 µL of the brain homogenate were then treated exactly as the plasma samples described above.

#### Standards and quality control (QC) samples

K_2_EDTA Blank mouse plasma purchased from Valley Biomedical, California, USA was used to prepare standards and QC samples for plasma quantitation and as surrogates for brain homogenate quantitation. Calibration samples ranged from 2.34 ng/mL to 4,800 ng/mL. QC samples concentration included 14 ng/mL (QC-L), 255 ng/mL (QC-M) and 3,600 ng/mL (QC-H). Standards and QC samples were processed the same way as the sample extracts described above.

### Analytical methods and statistics for plasma and tissue samples

Samples were analyzed by UHPLC-ESI MS/MS using a TQ-5500 Sciex triple quadrupole mass spectrometer equipped with a Shimadzu Nexera UHPLC pump and auto-sampler system using an ACE C18 PFP, 2.50 x 50 mm, 1.7 µ particle size column and gradient elution consisting of solvent A (0.1% formic acid in water) and solvent B (0.1% formic acid in acetonitrile) starting at 20% B from 0 min to 0.4 min and then increased to 100% B from 0.4 min to 0.6 min. At 2.0 min, the mobile phase composition was switched back to 60% B for 1 min. The flow rate used throughout the experiment was 0.4 min/mL. The analyte, NBI-921352, and the IS were detected by electrospray in the positive ion mode using the following transitions: m/z 460/91 for NBI-921352 and m/z 503/341m/z for the IS. The UHPLC-ESI MS/MS system was controlled by Analyst 1.6.

Sample concentrations were determined using a linear calibration function, weighted 1/X, generated by the regression of analyte to IS peak area ratios in the standard samples to their respective concentrations. Acceptance criteria for the analytical run required that the back calculated values of the standards and the QC samples fell within ± 20% of their nominal values, except for the lowest standard or lower limit of quantitation (LLOQ), for which the acceptance criterion was ± 25%. At least 6 out of 12 standard points had to show back-calculated values within ± 20% of their nominal concentrations for the calibration to be accepted. At least three QC samples, one at each level, had to show back-calculated values within ± 20% of their nominal concentrations for the whole sample batch to be valid.

### Animals

After delivery, animals were allowed sufficient time to acclimate prior to testing (∼1 week). All animals were housed in plastic cages in rooms with controlled humidity, ventilation, and lighting (12 hr/12 hr light–dark cycle). All animal procedures were performed using protocols approved by Xenon Animal Care Committee and the Canadian Council on Animal Care.

### Scn8aN1768D/+ mice

Xenon Pharmaceuticals Inc. licensed the mouse with the missense mutation p.Asn1768Asp (N1768D) in the neuronal sodium channel Na_V_1.6, characterized and developed by Dr. M Meisler (University Of Michigan, MI, USA). The *Scn8a^N1768D^* knock-in allele was generated by TALEN targeting of (C57BL/6JXSJL) F2 eggs at the University of Michigan Transgenic Animal Model Core. The line was propagated by backcrossing N1768D/+ heterozygotes to C57BL/6J wild-type mice (The Jackson Laboratory, Bar Harbor, ME). Male N1768D/+ heterozygotes on a C57BL/6J background were subsequently backcrossed to C3HeB/FeJ female mice. All the experiments were performed using animals following at least 7 such backcrosses. Experiments were performed using (B6 × C3He) F7 (F7. N1768D/+) offspring aged 35-42 days.

### WT mice

Adult male CF-1 WT albino mice 26-35 g were obtained from Charles River, Senneville, Quebec, Canada. All the assays were carried out in mice 9 weeks or older.

### Sprague-Dawley rats

Adult male Sprague-Dawley albino rats weighing 150-200 g were obtained from Envigo, Livermore, CA, USA. All the assays were carried out in rats 5 weeks or older.

### The modified 6 Hz psychomotor seizure assay

The modified 6 Hz seizure assay in *Scn8a^N1768D/+^* heterozygous mice was adapted from the traditional 6 Hz assay psychomotor seizure assay to provide a measure of in vivo on target (Na_V_1.6 mediated) efficacy (Barton et al., 2001). The modified assay used a low frequency (6 Hz) but long-duration stimulation (3 seconds) to induce seizures. We identified 12 mA and a 0.3 millisecond pulse interval as a suitable current for testing in *Scn8a^N1768D/+^* mice, since it differentiated mutant and wild-type (WT) mice. An electroshock (6 Hz, 12 mA) was delivered for 3 seconds (at 0.3 millisecond pulse interval) by corneal electrodes (Electro Convulsive Therapy Unit 57800 from Ugo Basile). Immediately prior to the electroshock, the animals’ eyes were anesthetized with a drop of Alcaine (0.5% proparacaine hydrochloride). Upon corneal stimulation, WT mice experienced mild seizure behaviors such as facial clonus, forelimb clonus, Straub tail, rearing, and falling, but did not experience a generalized tonic-clonic seizure (GTC) with hindlimb extension. *Scn8a^N1768D/+^* animals, however, in addition to mild seizure behaviors, experienced a GTC with hindlimb extension. The modified assay showed a clear differentiation of seizure behavior between WT and *Scn8a^N1768D/+^* mice. *Scn8a^N1768D/+^* mice exhibited GTC with hindlimb extension but not WT mice.

For the single-dose and repeated-dose efficacy experiments, *Scn8a^N1768D/+^* animals were dosed PO with vehicle or NBI-921352 two hours before the administration of the electric stimulation. An animal was considered protected in the assay upon prevention of GTC with hindlimb extension and was then scored “0”. An animal displaying GTC with hindlimb extension was considered not protected and is then scored “1”. The experimenter scoring the seizure behavior was blinded to the treatment.

### DC-Maximal electroshock seizure assay in rodents

The maximal electroshock seizure (MES) assay has been extensively used in the search for anticonvulsant substances (Loscher et al., 1991; Piredda et al., 1985; White et al., 1995). The MES assay is sensitive to nonselective NaV inhibitors. It is considered a model for generalized tonic-clonic (GTC) seizures and provides an assessment of seizure spread. Briefly, an electroshock of direct current (DC) was delivered by corneal electrodes (Electro Convulsive Therapy Unit 57800 from Ugo Basile). The parameters of stimulation were different between mice and rats. In CF1 mice, a direct current of 50 mA (60 Hz) was delivered for 0.2 seconds (pulse width of 0.5 ms), whereas in Sprague Dawley (SD) rats, a direct current of 150 mA (60 Hz) was delivered for 0.3 seconds (pulse width of 0.5 ms). Immediately prior to the electroshock, the animals’ eyes were anesthetized with a drop of Alcaine (0.5% proparacaine hydrochloride). Upon corneal stimulation, naïve animals experienced a generalized tonic-clonic seizure (GTC) with hindlimb extension.

For the efficacy experiments, single dose and repeated dose, animals were dosed PO with vehicle or NBI-921352 two hours before the administration of the electric stimulation. An animal was considered protected in the assay in the absence of a GTC with hindlimb extension and is then scored “0”. An animal displaying GTC with hindlimb extension was considered not protected and is then scored “1”. The experimenter scoring the seizure behavior was blinded to the treatment.

### Blinding of in vivo efficacy experiments

On each testing day, individual treatment groups were assigned a random label (e.g., A, B, C, etc.) by the technical staff administering the compound. To ensure blinding, the technical staff member performing drug administration differed from the person performing the test. Therefore, the experimenter conducting testing was blinded to treatment group (e.g., drug or vehicle treatment, dose, and time point).

### Randomization of in vivo efficacy experiments

Randomization of animals into various treatment groups occurred on a per-animal (e.g., rather than a per-cage) basis. Therefore, each animal was randomly assigned to a treatment group, and all animals tested in each experiment had an equal chance of assignment to any treatment group. Prior to each study, a randomization sequence was obtained (www.graphpad.com/quickcalcs).

## Acknowledgements

We thank Dr. Fiona Scott, PhD of Neurocrine Biosciences for thoughtful discussions and commentary on this manuscript. We thank Ian Mortimer of Xenon Pharmaceuticals for strategic discussions. We thank Dr. Miriam Meisler, PhD for making *Scn8a^N1768D/+^* mice available. We thank the *SCN8A*-RES patients, families and patient advocates for detailed discussions regarding the clinical presentation of *SCN8A* mutations and unmet medical needs in their community.

## References

Ahuja, S., Mukund, S., Deng, L., Khakh, K., Chang, E., Ho, H., … Payandeh, J. (2015). Structural basis of Nav1.7 inhibition by an isoform-selective small-molecule antagonist. Science, 350(6267), aac5464. doi:10.1126/science.aac5464

Barbosa, C., Tan, Z. Y., Wang, R., Xie, W., Strong, J. A., Patel, R. R., … Cummins, T. R. (2015). Navbeta4 regulates fast resurgent sodium currents and excitability in sensory neurons. Mol Pain, 11, 60. doi:10.1186/s12990-015-0063-9

Barton, M. E., Klein, B. D., Wolf, H. H., & White, H. S. (2001). Pharmacological characterization of the 6 Hz psychomotor seizure model of partial epilepsy. Epilepsy Res, 47(3), 217–227. doi:10.1016/s0920-1211(01)00302-3

Bean, B. P., Cohen, C. J., & Tsien, R. W. (1983). Lidocaine block of cardiac sodium channels. J Gen Physiol, 81(5), 613–642. doi:10.1085/jgp.81.5.613

Ben-Shalom, R., Keeshen, C. M., Berrios, K. N., An, J. Y., Sanders, S. J., & Bender, K. J. (2017). Opposing Effects on NaV1.2 Function Underlie Differences Between SCN2A Variants Observed in Individuals With Autism Spectrum Disorder or Infantile Seizures. Biol Psychiatry, 82(3), 224–232. doi:10.1016/j.biopsych.2017.01.009

Boerma, R. S., Braun, K. P., van de Broek, M. P., van Berkestijn, F. M., Swinkels, M. E., Hagebeuk, E. O., … Koeleman, B. P. (2016). Remarkable Phenytoin Sensitivity in 4 Children with SCN8A-related Epilepsy: A Molecular Neuropharmacological Approach. Neurotherapeutics, 13(1), 192–197. doi:10.1007/s13311-015-0372-8

Braakman, H. M., Verhoeven, J. S., Erasmus, C. E., Haaxma, C. A., Willemsen, M. H., & Schelhaas, H. J. (2017). Phenytoin as a last-resort treatment in SCN8A encephalopathy. Epilepsia Open, 2(3), 343–344. doi:10.1002/epi4.12059

Burgess, D. L., Kohrman, D. C., Galt, J., Plummer, N. W., Jones, J. M., Spear, B., & Meisler, M. H. (1995). Mutation of a new sodium channel gene, Scn8a, in the mouse mutant ’motor endplate disease’. Nat Genet, 10(4), 461–465. doi:10.1038/ng0895-461

Caron, J., & Libersa, C. (1997). Adverse effects of class I antiarrhythmic drugs. Drug Saf, 17(1), 8–36. doi:10.2165/00002018-199717010-00002

Catterall, W. A., Kalume, F., & Oakley, J. C. (2010). NaV1.1 channels and epilepsy. J Physiol, 588(Pt 11), 1849–1859. doi:10.1113/jphysiol.2010.187484

Chen, Q., Kirsch, G. E., Zhang, D., Brugada, R., Brugada, J., Brugada, P., … Wang, Q. (1998). Genetic basis and molecular mechanism for idiopathic ventricular fibrillation. Nature, 392(6673), 293–296. doi:10.1038/32675

Claes, L., Del-Favero, J., Ceulemans, B., Lagae, L., Van Broeckhoven, C., & De Jonghe, P. (2001). De novo mutations in the sodium-channel gene SCN1A cause severe myoclonic epilepsy of infancy. Am J Hum Genet, 68(6), 1327–1332. doi:10.1086/320609

Courtney, K. R., Kendig, J. J., & Cohen, E. N. (1978). The rates of interaction of local anesthetics with sodium channels in nerve. J Pharmacol Exp Ther, 207(2), 594–604. Retrieved from https://www.ncbi.nlm.nih.gov/pubmed/712641

Du, J., Simmons, S., Brunklaus, A., Adiconis, X., Hession, C. C., Fu, Z., … Lal, D. (2020). Differential excitatory vs inhibitory SCN expression at single cell level regulates brain sodium channel function in neurodevelopmental disorders. Eur J Paediatr Neurol, 24, 129–133. doi:10.1016/j.ejpn.2019.12.019

Encinas, A. C., Watkins, J. C., Longoria, I. A., Johnson, J. P., Jr., & Hammer, M. F. (2020). Variable patterns of mutation density among NaV1.1, NaV1.2 and NaV1.6 point to channel-specific functional differences associated with childhood epilepsy. PLoS One, 15(8), e0238121. doi:10.1371/journal.pone.0238121

Escayg, A., MacDonald, B. T., Meisler, M. H., Baulac, S., Huberfeld, G., An-Gourfinkel, I., … Malafosse, A. (2000). Mutations of SCN1A, encoding a neuronal sodium channel, in two families with GEFS+2. Nat Genet, 24(4), 343–345. doi:10.1038/74159

FDA. (2021). Drug Safety Communication: Studies show increased risk of heart rhythm problems with seizure and mental health medicine lamotrigine (Lamictal) in patients with heart disease. Retrieved from https://www.fda.gov/drugs/drug-safety-and-availability/studies-show-increased-risk-heart-rhythm-problems-seizure-and-mental-health-medicine-lamotrigine

Focken, T., Burford, K., Grimwood, M. E., Zenova, A., Andrez, J. C., Gong, W., … Empfield, J. R. (2019). Identification of CNS-Penetrant Aryl Sulfonamides as Isoform-Selective NaV1.6 Inhibitors with Efficacy in Mouse Models of Epilepsy. J Med Chem, 62(21), 9618–9641. doi:10.1021/acs.jmedchem.9b01032

Gardella, E., & Moller, R. S. (2019). Phenotypic and genetic spectrum of SCN8A-related disorders, treatment options, and outcomes. Epilepsia, 60 *Suppl 3*, S77–S85. doi:10.1111/epi.16319

Gennaro, E., Veggiotti, P., Malacarne, M., Madia, F., Cecconi, M., Cardinali, S., … Zara, F. (2003). Familial severe myoclonic epilepsy of infancy: truncation of Nav1.1 and genetic heterogeneity. Epileptic Disord, 5(1), 21–25. Retrieved from https://www.ncbi.nlm.nih.gov/pubmed/12773292

Hammer, M. F., Wagnon, J. L., Mefford, H. C., & Meisler, M. H. (2016). SCN8A-Related Epilepsy with Encephalopathy. In M. P. Adam, H. H. Ardinger, R. A. Pagon, S. E. Wallace, L. J. H. Bean, K. Stephens, & A. Amemiya (Eds.), GeneReviews((R)). Seattle (WA).

Hawkins, N. A., Martin, M. S., Frankel, W. N., Kearney, J. A., & Escayg, A. (2011). Neuronal voltage-gated ion channels are genetic modifiers of generalized epilepsy with febrile seizures plus. Neurobiol Dis, 41(3), 655–660. doi:10.1016/j.nbd.2010.11.016

Inglis, G. A. S., Wong, J. C., Butler, K. M., Thelin, J. T., Mistretta, O. C., Wu, X., … Escayg, A. (2020). Mutations in the Scn8a DIIS4 voltage sensor reveal new distinctions among hypomorphic and null Nav 1.6 sodium channels. Genes Brain Behav, 19(4), e12612. doi:10.1111/gbb.12612

Johannesen, K. M., Gardella, E., Encinas, A. C., Lehesjoki, A. E., Linnankivi, T., Petersen, M. B., … Moller, R. S. (2019). The spectrum of intermediate SCN8A-related epilepsy. Epilepsia, 60(5), 830–844. doi:10.1111/epi.14705

Liu, Y., Schubert, J., Sonnenberg, L., Helbig, K. L., Hoei-Hansen, C. E., Koko, M., … Lerche, H. (2019). Neuronal mechanisms of mutations in SCN8A causing epilepsy or intellectual disability. Brain, 142(2), 376–390. doi:10.1093/brain/awy326

Loscher, W., Fassbender, C. P., & Nolting, B. (1991). The role of technical, biological and pharmacological factors in the laboratory evaluation of anticonvulsant drugs. II. Maximal electroshock seizure models. Epilepsy Res, 8(2), 79–94. doi:10.1016/0920-1211(91)90075-q

Martin, M. S., Tang, B., Papale, L. A., Yu, F. H., Catterall, W. A., & Escayg, A. (2007). The voltage-gated sodium channel Scn8a is a genetic modifier of severe myoclonic epilepsy of infancy. Hum Mol Genet, 16(23), 2892–2899. doi:10.1093/hmg/ddm248

Mason, E. R., Wu, F., Patel, R. R., Xiao, Y., Cannon, S. C., & Cummins, T. R. (2019). Resurgent and Gating Pore Currents Induced by De Novo SCN2A Epilepsy Mutations. eNeuro, 6(5). doi:10.1523/ENEURO.0141-19.2019

McCormack, K., Santos, S., Chapman, M. L., Krafte, D. S., Marron, B. E., West, C. W., … Castle, N. A. (2013). Voltage sensor interaction site for selective small molecule inhibitors of voltage-gated sodium channels. Proc Natl Acad Sci U S A, 110(29), E2724–2732. doi:10.1073/pnas.1220844110

Meisler, M. H. (2019). SCN8A encephalopathy: Mechanisms and models. Epilepsia, 60 *Suppl 3*, S86–S91. doi:10.1111/epi.14703

Mistry, A. M., Thompson, C. H., Miller, A. R., Vanoye, C. G., George, A. L., Jr., & Kearney, J. A. (2014). Strain- and age-dependent hippocampal neuron sodium currents correlate with epilepsy severity in Dravet syndrome mice. Neurobiol Dis, 65, 1–11. doi:10.1016/j.nbd.2014.01.006

Neurocrine. (2019). Neurocrine Biosciences and Xenon Pharmaceuticals Announce Agreement to Develop First-in-Class Treatments for Epilepsy. *Press Release*. Retrieved from https://neurocrine.gcs-web.com/news-releases/news-release-details/neurocrine-biosciences-and-xenon-pharmaceuticals-announce

Neurocrine. (2021). Study to Evaluate NBI-921352 as Adjunctive Therapy in Subjects With SCN8A Developmental and Epileptic Encephalopathy Syndrome (SCN8A-DEE). *ClinicalTrials.gov* Retrieved from https://www.clinicaltrials.gov/ct2/show/NCT04873869?term=NCT04873869&draw=2&rank=1

Pan, Y., & Cummins, T. R. (2020). Distinct functional alterations in SCN8A epilepsy mutant channels. J Physiol, 598(2), 381–401. doi:10.1113/JP278952

Piredda, S. G., Woodhead, J. H., & Swinyard, E. A. (1985). Effect of stimulus intensity on the profile of anticonvulsant activity of phenytoin, ethosuximide and valproate. J Pharmacol Exp Ther, 232(3), 741–745. Retrieved from https://www.ncbi.nlm.nih.gov/pubmed/3919174

Potet, F., Egecioglu, D. E., Burridge, P. W., & George, A. L., Jr. (2020). GS-967 and Eleclazine Block Sodium Channels in Human Induced Pluripotent Stem Cell-Derived Cardiomyocytes. Mol Pharmacol, 98(5), 540–547. doi:10.1124/molpharm.120.000048

Ptacek, L. J., George, A. L., Jr., Griggs, R. C., Tawil, R., Kallen, R. G., Barchi, R. L., … Leppert, M. F. (1991). Identification of a mutation in the gene causing hyperkalemic periodic paralysis. Cell, 67(5), 1021–1027. doi:10.1016/0092-8674(91)90374-8

Raman, I. M., & Bean, B. P. (1997). Resurgent sodium current and action potential formation in dissociated cerebellar Purkinje neurons. J Neurosci, 17(12), 4517–4526. Retrieved from https://www.ncbi.nlm.nih.gov/pubmed/9169512

Raman, I. M., Sprunger, L. K., Meisler, M. H., & Bean, B. P. (1997). Altered subthreshold sodium currents and disrupted firing patterns in Purkinje neurons of Scn8a mutant mice. Neuron, 19(4), 881–891. doi:10.1016/s0896-6273(00)80969-1

Rojas, C. V., Wang, J. Z., Schwartz, L. S., Hoffman, E. P., Powell, B. R., & Brown, R. H., Jr. (1991). A Met-to-Val mutation in the skeletal muscle Na+ channel alpha-subunit in hyperkalaemic periodic paralysis. Nature, 354(6352), 387–389. doi:10.1038/354387a0

Royeck, M., Horstmann, M. T., Remy, S., Reitze, M., Yaari, Y., & Beck, H. (2008). Role of axonal NaV1.6 sodium channels in action potential initiation of CA1 pyramidal neurons. J Neurophysiol, 100(4), 2361–2380. doi:10.1152/jn.90332.2008

Strichartz, G. (1976). Molecular mechanisms of nerve block by local anesthetics. Anesthesiology, 45(4), 421–441. doi:10.1097/00000542-197610000-00012

Tidball, A. M., Lopez-Santiago, L. F., Yuan, Y., Glenn, T. W., Margolis, J. L., Clayton Walker, J., … Parent, J. M. (2020). Variant-specific changes in persistent or resurgent sodium current in SCN8A-related epilepsy patient-derived neurons. Brain, 143(10), 3025–3040. doi:10.1093/brain/awaa247

Veeramah, K. R., O’Brien, J. E., Meisler, M. H., Cheng, X., Dib-Hajj, S. D., Waxman, S. G., … Hammer, M. F. (2012). De novo pathogenic SCN8A mutation identified by whole-genome sequencing of a family quartet affected by infantile epileptic encephalopathy and SUDEP. Am J Hum Genet, 90(3), 502–510. doi:10.1016/j.ajhg.2012.01.006

Wagnon, J. L., Korn, M. J., Parent, R., Tarpey, T. A., Jones, J. M., Hammer, M. F., … Meisler, M. H. (2015). Convulsive seizures and SUDEP in a mouse model of SCN8A epileptic encephalopathy. Hum Mol Genet, 24(2), 506–515. doi:10.1093/hmg/ddu470

Wagnon, J. L., & Meisler, M. H. (2015). Recurrent and Non-Recurrent Mutations of SCN8A in Epileptic Encephalopathy. Front Neurol, 6, 104. doi:10.3389/fneur.2015.00104

Wengert, E. R., & Patel, M. K. (2021). The Role of the Persistent Sodium Current in Epilepsy. Epilepsy Curr, 21(1), 40–47. doi:10.1177/1535759720973978

Wengert, E. R., Tronhjem, C. E., Wagnon, J. L., Johannesen, K. M., Petit, H., Krey, I., … Moller, R. S. (2019). Biallelic inherited SCN8A variants, a rare cause of SCN8A-related developmental and epileptic encephalopathy. Epilepsia, 60(11), 2277–2285. doi:10.1111/epi.16371

White, H. S., Johnson, M., Wolf, H. H., & Kupferberg, H. J. (1995). The early identification of anticonvulsant activity: role of the maximal electroshock and subcutaneous pentylenetetrazol seizure models. Ital J Neurol Sci, 16(1-2), 73–77. doi:10.1007/BF02229077

Yu, F. H., Mantegazza, M., Westenbroek, R. E., Robbins, C. A., Kalume, F., Burton, K. A., … Catterall, W. A. (2006). Reduced sodium current in GABAergic interneurons in a mouse model of severe myoclonic epilepsy in infancy. Nat Neurosci, 9(9), 1142–1149. doi:10.1038/nn1754

Zaman, T., Abou Tayoun, A., & Goldberg, E. M. (2019). A single-center SCN8A-related epilepsy cohort: clinical, genetic, and physiologic characterization. Ann Clin Transl Neurol, 6(8), 1445–1455. doi:10.1002/acn3.50839

